# Signals of positive selection in Palearctic bat species coexisting with a fungal pathogen

**DOI:** 10.1101/2023.12.04.569365

**Authors:** VG Twort, Veronika N Laine, K Field, F Whiting-Fawcett, F Ito, TM Lilley

## Abstract

Traits that directly influence the survival of an organism are suspect to positive selection. Disease can act as a driving force in shaping the genetic makeup across populations, even species, if the impacts are influencing a particularly sensitive part of their life cycles. White-nose syndrome is a fungal disease that affects bats during hibernation. The mycosis has caused massive population declines of susceptible species in North America, whereas in Eurasia, where the fungal pathogen has coevolved with its hosts for an extended period of time, bats appear to tolerate infection. Here, we adopted both whole-genome sequencing approaches and a literature search to compile a set of 300 genes from which to investigate for signals of positive selection in genomes of 11 Eurasian bats at the codon-level. Our results indicate significant positive selection in 38 genes, many of which have a marked role in responses to infection. Our findings suggest the fungal disease known as white-nose syndrome may have applied a significant selective pressure on hibernatory Eurasian Myotis-bats in the past, which can partially explain their survival in the presence of the pathogen.

## Introduction

Infectious diseases are natural processes affecting wildlife and contribute to ecosystem stability (Cunningham et al. 2017). However, over the past decades, these processes have been intensified due to anthropogenic activities, such as urbanisation, and environmental and climate change (Baylis & Risley 2012, Cunningham et al. 2017, Baker et al. 2022). Furthermore, the anthropogenic impacts on the incidence of epizootic diseases have been identified as an increasing threat to wildlife conservation (Cunningham et al. 2017, Baker et al. 2022). Recently, the role of infectious diseases in population declines and/or extinction has received a growing recognition, but there is still a critical need to identify and understand the impacts and projections of diseases on populations (Daszak et al. 2000).

White-nose syndrome (WNS) is a wildlife disease caused by a fungal infection affecting hibernating bats in North America (Blehert et al. 2009). It is considered one of the most detrimental wildlife diseases of recent decades (Frick et al. 2016), with mass mortality of affected species causing unprecedented population collapses in many of the affected areas in North America (Blehert et al. 2009, Frick et al. 2015). The causative agent, the cold-loving fungus *Pseudogymnoascus destructans* (=*Geomyces destructans*, (Blehert et al. 2009), is endemic to the Palearctic, where it does not cause significant mortality in bat hosts, due to extended coevolution between the pathogen and the hosts (Fischer et al. 2020, Whiting-Fawcett et al. 2021).

Hibernation allows bats to survive periods of food scarcity by reducing their energy expenditure via decreased metabolism. Host mortality caused by WNS occurs via the disruption of the normal pattern of hibernation (Reeder et al. 2012, Warnecke et al. 2012). Infected North American *Myotis lucifugus* arouse three times more frequently in the final third of the hibernation period (Warnecke et al. 2012), expending large amounts of the fat reserves that are expected to last until insect food is available again in the spring. The fungus invades host tissue during the extended bouts of torpor when the host immune system is downregulated (Meteyer et al. 2009). *Pseudogymnoascus destructans* infection induces the production of inflammatory cytokines during the arousals that take place during hibernation (Field et al. 2015, 2018, Lilley et al. 2017). The irritation, such as pain and itchiness, associated with inflammation, in addition to compounds produced by the fungus (Flieger et al. 2016) and evaporative water loss from open sores (McGuire et al. 2017) has been suggested to increase the frequency of arousals. This leads to emaciation and eventual death in the more susceptible bat species.

In the Palearctic, *P. destructans* infects a number of different bat species, mostly in the genus *Myotis* (Hoyt et al. 2015, Zukal et al. 2016). The coevolution between the fungus and the bat hosts is extensive (Leopardi et al. 2015, Fischer et al. 2020), and although no direct evidence exists, it has been suggested that *Myotis* species may have experienced similar population declines as a result of disease in the past (Martinkova et al. 2010). The most extensively studied Palearctic species, *Myotis myotis*, elicits no response to infection (Fritze et al. 2019, 2021, Lilley et al. 2019), which suggests it utilises tolerance as a response to infection (Ayres & Schneider 2012). Tolerance allows the host to evade harmful immunopathology that appears to be a major factor contributing to mortality of bats in the Nearctic.

Diseases, such as WNS, are a strong driving force of natural selection (Karlsson et al. 2014). Selection pressures result in the modification of the hosts’ genetic diversity and leave behind distinctive signatures in the genome. The nature of these signatures depends on the evolutionary timescale of interest. Among these identifiable footprints of selection are selective sweeps, whereby an advantageous mutation eliminates or reduces variation in the population at linked neutral sites as the mutation increases in frequency. Alternatively, alterations in host genetic makeup can be detected in protein coding regions as signals of natural selection. Whereby, the rates of nucleotide changes that either change (non-synonymous or d_N_) or preserve (synonymous or d_S_) the protein sequence are used to infer selection. Under neutral evolution the rates are equal (d_N_/d_S_ = 1). Whereas, an excess of non-synonymous changes is a sign of positive selection (d_N_/d_S_ > 1), while the converse is associated with a predominant negative selection pressure (d_N_/d_S_ < 1) (Yang & Bielawski 2000). Based on the idea of a long term association between Palearctic bats and *P. destructans* (Leopardi et al. 2015), it is likely that bats have evolved inheritable mechanisms leading to infection tolerance (Whiting-Fawcett et al. 2021). Signatures of such mechanisms could be detected within protein coding genes associated with wound healing, immune responses and hibernation physiology (Field et al. 2015, Harazim et al. 2018a, Lilley et al. 2019).

Because the fungus causing WNS, *P. destructans*, is endemic to the Palearctic (Leopardi et al. 2015), but infected bat hosts do not experience significant mortality (Puechmaille et al. 2011) or elicit a transcriptional response to infection (Lilley et al. 2019), we hypothesise that Palearctic species susceptible to infection show signs of positive selection in genes that are associated with immune system signalling and function. To investigate the impact of WNS on the hosts’ genetic makeup, we use a variety of approaches to detect selection. First, we constructed a whole-genome dataset of 12 *Myotis* species (1 Nearctic, 11 Palearctic) to investigate the general overall patterns of selection across the phylogeny and conduct a branch test to detect any lineage specific selection. We also will detect recent positive selection with selective sweep analysis using single nucleotide polymorphisms (SNPs) in *Myotis myotis* and *Myotis lucifugus* separately. We also conduct a systematic literature search to list genes already found to be associated with host responses during infection with *P. destructans*. Finally, the combined gene set curated from a literature search, sweep analysis and the whole-genome selection patterns is used to test for the presence of inheritable variation within *Myotis*. The identification of positive selection among these genes may highlight variations that contribute to tolerance among Palearctic lineages of *Myotis*. The identified gene variants and mechanisms contributing to survival in the Palearctic can be applied to conservation genetic approaches in North America to facilitate more effective management methods.

## Methods

### 1. Sample Collection, DNA Extraction and Sequencing

Samples for DNA extraction were obtained from existing collections of the authors collected under relevant permits in their respective countries. No bats were sampled as a part of the present study.

Eleven out of the twelve species sampled represent Myotis from the Palearctic clade (*M. bechsteinii*, *M. daubentonii*, *M. alcathoe*, *M. mystacinus*, *M. dasycneme*, *M. frater*, *M. pequinus*, *M. petax*, *M. myotis*, *M. nattereri*, *M. ikonnikovi*) with distributions ranging from the Iberian Peninsula to Japan. All species reside in the temperate or boreal zone and utilise extended periods of torpor, hibernation, over the winter months (Wilson & Mittermeier 2019). The final species is *M. lucifugus*, the only representative of the Nearctic clade of Myotis in our study. The species has been extensively studied with regards to WNS (Reeder et al. 2012, Warnecke et al. 2012, Cheng et al. 2019, Frick et al. 2022) and its responses to infection have been documented (Field et al. 2015, 2018, Lilley et al. 2017, 2019). The purpose of the species in our data set is to act as a comparison to which Palearctic species are compared against. Full details for each sample are given in Supplementary Table 1.

DNA extraction was carried out using one of the following methods: 1) QIAmp DNA Mini Kit (Qiagen, Germany); 2) DNeasy Blood and Tissue Kit (Qiagen, Germany); 3) SDS Extraction method (full details can be found in Supplementary Methods 1), with extractions being stored at -80 °C. Library preparation and sequencing was carried out by a variety of providers (see Supplementary Table 1) as follows. For samples processed at: 1) Novogene, United Kingdom: Library preparation was carried out with the Novogene NGS DNA Library Preparation Set (Cat No. PT004) prior to sequencing on a NovaSeq6000 platform to generate 2 x 150 bp reads. 2) Novogene, China: 1.5 µg of DNA was used as input for the TruSeq Nano DNA HT Sample preparation kit (Illumina), with a 350 bp insert. Samples were sequenced on a NovaSeq6000 to generate 2 x 150 bp reads. 3) DNA Sequencing and Genomics Laboratory (BIDGEN): Libraries were prepared using the Nextera™ DNA Flex Library Preparation Kit (Illumina) and sequenced on a NovaSeq6000 to generate 2 x 150 bp reads. 4) University of Liverpool Centre for Genomics Research (CGR): Libraries were prepared using the NEBNext Ultra II FS Kit (∼300 bp inserts) on the Mosquito platform using a 1/10 reduced volume protocol. Samples were sequenced on a NovaSeq6000 to generate 2 x 150 bp reads. For those samples sequenced at CGR, the provider also carried out read trimming as follows: Adapter removal with Cutadapt (version 1.2.1, (Martin 2011), followed by trimming with Sickle (version 1.200 (Joshi & Fass 2011)), with a minimum window quality score of 20, and removal of reads less than 15 bp.

All raw sequencing reads have been submitted to the NCBI SRA under bioproject XXXX. In addition to the 18 samples sequenced here, sequencing data from a further six *M. lucifugus*, originally sampled at the onset of the WNS-epizootic in North America (Lilley et al. 2020), were downloaded from the NCBI SRA (see Supplementary Table 2).

### 2. Data Processing and Read Mapping

Raw reads were quality checked with FASTQC (version 0.11.8, (Andrews 2015), followed by ambiguous base removal with Prinseq (version 0.20.4, (Schmieder & Edwards 2011). Adapter removal and quality filtering was undertaken with Trimmomatic (version 0.39, (Bolger et al. 2014), using library appropriate adapter sequences and the following settings: ILLUMINACLIP: :2:30:10 LEADING:3 TRAILING:3 SLIDINGWINDOW:4:25 MINLEN:50.

All samples were mapped against the *M. myotis* genome (GCF_014108235.1 mMyoMyo1.p) using bowtie2 (version 2.4.1 (Langmead & Salzberg 2012)) utilising the sensitive-local option. The resulting sam file was converted to a sorted bam file with samtools (version 1.10 (Danecek et al. 2021)). Duplicate removal was carried out with Picard (version 2.27.5 (“Picard Tools ȓ By Broad Institute”)) MarkDuplicates option, with a lenient validation stringency, coordinate sort order and the removal of duplicates. Appropriate reads groups were added to each sample using the AddorReplaceReadGroups function of Picard, with the resulting bam file was indexed with samtools. A consensus assembly was created for each sample with ANGSD (version 0.939, (Korneliussen et al. 2014) using the following settings: -minQ 20 -doCounts 1 -minMapQ 20 -setMinDepth 3 -dofasta 2.

### 3. Phylogenetic Dataset Construction and Analysis

#### 3.1 Orthologue Identification

In order to identify single copy orthologues, the BUSCO Mammalia_odb10 reference dataset (Manni et al. 2021) was used. At the time of analysis, the current version of BUSCO did not output nucleotide sequences, therefore Metaeuk (Release 5-34c21f2, (Levy Karin et al. 2020)) was used to identify gene sequences (settings: easy-predict –cov 0.6 –filter-mas 1 –metaeuk-eval 0.0001 -metaeuk-tcov 0.6 –min-length 40). Result files were filtered to remove any duplicated genes and genes identified in less than 8 individuals. Both the protein and nucleotide sequences produced by Metaeuk were taken forward.

Initial protein alignments were generated using MAFFT (version 7.429, (Katoh & Standley 2013)), using the ‘auto’ option and manually checked to ensure accuracy, screened for the presence of pseudogenes, reading frame errors and alignment errors using Geneious 11.0.3 (https://www.geneious.com). After screening, any alignments with less than 8 sequences, <50% of the ORF or those lacking either *M. myotis* or *M. lucifugus* sequences were discarded. This resulted in a final dataset of 2,515 genes (Supplementary Table 3). Amino acid alignments were converted into codon alignments utilising the nucleotide sequences extracted by Metaeuk with Pal2Nal (Version 14, (Suyama et al. 2006)). Alignments were cleaned with gblocks (0.91b, (Castresana 2000)) with the following settings: -t=c, -b3=8, -b5=15 -b5=a, b1 = 50% of the number of sequences +2 and b2 = 50% of the number of sequences +4. The final gene alignments can be accessed at Zenodo DOI: XXXXX. Annotations for each BUSCO gene code were retrieved from OrthoDB v10 (Kriventseva et al. 2019).

#### 3.2 Phylogenetic Analysis

To reconstruct the evolutionary relationships a multispecies coalescent analysis was used. Individual gene trees were reconstructed with IQ-TREE2 (2.1.3, (Minh et al. 2020)), with ModelFinder (Kalyaanamoorthy et al. 2017) being run to determine the optimal model and 1000 ultrafast bootstraps (UFBoot2) approximations (Hoang et al. 2018). For species tree inference two approaches were undertaken with ATSTRAL-III (5.7.8, (Zhang et al. 2018)), first default parameters were used, while the second analysis incorporated the bootstrap trees from IQ-TREE2 with 500 replicates.

#### 3.3 Inferring Selection Associated with *M. myotis* and *M. lucifugus*

Branch tests were implemented to detect selection acting on particular lineages. Two independent analyses were carried out, in the first analysis the branch leading to *Myotis myotis* was labelled as the foreground branch, while in the second the branch leading to *M. lucifugu*s was considered the foreground. For each analysis individual gene trees were used. In order for the labelled branches to be read correctly, all of the extra information associated with each branch, such as the branch length and bootstrap scores, were removed with PareTree (version 1.02, Available from: http://emmahodcroft.com/PareTree.html), followed by the branch of interest being labelled. Branch models allow for the labelled (or foreground) branch to have an Omega (ω) independent of the rest of the phylogeny and were carried out with Paml 4.9j (Yang 2007). P-values were generated from the LRT tests using the R software (version 4.1.2, (R Core Team 2022)) function pchisq, with correction for multiple tests being carried out with the holm-bonferroni correction using the p.adjust function.

### 4. SNP Calling, Selective Sweeps Detection and Nucleotide Diversity

SNPs were called from *Myotis myotis* and *Myotis lucifugus*, seven individuals from each species, for selective sweep detection. Because of low numbers of individuals and low sequencing coverage in many of the individuals, we used two SNP callers after which overlapping SNPs were extracted. For *Myotis myotis* we used the aligned deduplicated files with read groups from section 2 for both callers. We also aligned *Myotis lucifugus* samples to a draft chromosome-level *Myotis lucifugus* assembly (mMyoLuc1, NCBI BioProject PRJNA973719) using the same settings as the *M. myotis* samples (see section 2), with deduplication and read group addition. The first caller was ANGSD version 0.940 (Kim et al. 2011, Korneliussen et al. 2014) with settings -uniqueOnly 1 -remove_bads 1 – only_proper_pairs 1 -C 50 -baq 1 -minMapQ 20 -minQ 20 -minInd 7 -doCounts 1 -setMinDepth 10 -setMaxDepth 100 -GL 2 -doMajorMinor 1 -doMaf 1 -SNP_pval 1e-6 -doPost 2 -doGeno 3 -doBcf 1 -doGlf 2. The second SNP caller was GATK HaplotypeCaller version 4.3.0.0 (Korneliussen et al. 2014) with -ERC GVCF and by scaffold (-L). The scaffolds were combined with picard’s GatherVcfs function. All the individual vcfs were combined with GATK’s CombineGVCFs tool. The variants were genotyped with GATK GenotypeGVCFs with settings --heterozygosity 0.001 and -stand-call-conf 20 and SNPs selected with GATK SelectVariants with settings --select-type-to-include SNP --restrict-alleles-to BIALLELIC. SNPs were filtered GATK’s VariantFiltration with --filter-expression “QD < 2.0 || FS > 60.0 || MQ < 40.0 || MappingQualityRankSum < -12.5 || SOR > 3.00 || ReadPosRankSum > -8.00” --filter-name “snp_filter” and selected from the files with vcftools --remove-filtered-all, version 0.1.16 (Danecek et al. 2011).

Both SNP caller files from both species were filtered with vcftools using settings --min-alleles 2 --max-alleles 2 --minDP 5 --maxDP 100 --max-missing 1. In order to select the overlapping SNPs from the files, we used Bedtools intersect version 2.30.0 (Quinlan & Hall 2010) (bedtools intersect -u -a GATK.vcf -b ANGSD.vcf) and filtered the overlapping SNPs with vcftools -- positions. Finally, the SNPs with differing genotypes within an individual from the overlap set was first detected with vcftools --diff-site-discordance and removed with vcftools --exclude-positions.

For the selective sweep detection we used RAiSD version 2.9 (Alachiotis & Pavlidis 2018) excluding known genome gaps. RAiSD was run for each species and the resulting outlier windows (highest μ values 0.05%) were filtered using a conservative threshold (α = 0.0005) in R version 4.3.0. The genes from these windows that were above the threshold were extracted with bedtools intersect using *Myotis myotis* NCBI gene annotation release 100 version GCF_014108235.1_mMyoMyo1 in *Myotis myotis*. In *Myotis lucifugus* samples a draft annotation was used. The mMyoLuc1 draft assembly was annotated by porting over gene annotations from mMyoMyo1 using Liftoff (v1.6.3, (Shumate & Salzberg 2021) and the settings “-exclude_partial -cds -polish;” these annotations were further cleaned using AGAT (v0.9.2, (Dainat et al. 2023).

Nucleotide diversity, Theta Watterson, and Tajima’s D were estimated for both species separately using ANGSD. First, the dosaf 1 function was used to calculate the site allele frequency spectrum likelihood (saf) for each species based on individual genotype likelihoods using the same quality specification as in genotype calling. Then, the realSFS function was used to optimize the saf and estimate the unfolded site frequency spectrum (SFS; Nielsen et al. 2012). Nucleotide diversity, Theta Watterson, and Tajima’s D were calculated for each site with the commands saf2theta and thetaStat in ANGSD.

### 5. Curated Gene Dataset

#### 5.1 Literature Search

We carried out a literature search to identify genes previously determined to be linked to White-Nose tolerance, using Web of Science (Clarivate) (carried out on 17/04/2023). The presence of the following keywords were searched for in the abstract or title or keywords: ((gene OR genome OR genetic OR transcript*) AND (*Pseudogymanoascus destructans* OR WNS OR white-nose syndrome OR white-nose disease)). The search yielded 134 papers. Upon closer investigation, 11 of these papers looked at bat genetic changes associated with WNS, yielding a total of 172 genes (Supplementary Table 4A).

In addition to the genes identified in the literature search, the genes identified in the sweep analysis (see Supplementary Table 4B) and the genes with the top 1% of ω values (from the phylogenetic dataset, see section 3.3 and Supplementary Table 4C) were also included. Gene codes, and aliases were checked against GeneCards (Stelzer et al. 2016). This resulted in a total of 347 genes of interest (Supplementary Table 4D).

#### 5.2 Gene Identification, Extraction and Alignment

To extract the coding (CDS) region of each gene, the *Myotis myotis* NCBI gene annotation release 100 version GCF_014108235.1_mMyoMyo1 was searched for corresponding CDS entries. In the case where genes had multiple isoforms, the longest version was extracted. Exon extraction from the mMyoMyo1 genome was carried out using bedtools getfasta (v2.30.0 (Quinlan & Hall 2010)) with the force strandedness option, followed by concatenation to create a reference CDS sequence. To ensure correct assembly of each gene, a blastp search was carried out against the NCBI nr database. These assembled CDS sequences were used as a reference set, to ensure correct extraction and assembly of sample CDS regions.

Exons were extracted from each consensus assembly using bedtools getfasta using the force strandedness option and assembled against the reference sequence using the ‘assemble to reference’ option in Geneious, with the subsequent CDS assemblies being aligned using MAFFT, using the auto option. Alignments were manually screened for frameshifts and difficult to align regions, those with less than 8 sequences, <80% of the CDS region and those lacking sequences from *M. lucifugus* and *M. myotis* were discarded. This resulted in a final dataset of 300 genes (Supplementary Table 5). Alignments were automatically cleaned with gblocks with the same settings as in section 3.1. The final gene alignments can be accessed from Zenodo DOI: XXXXX.

#### 5.3 Site Selection Patterns

To detect patterns of selection within individual genes, site-based likelihood models were used. For each gene, individual gene trees were first constructed under the maximum likelihood framework with IQ-TREE2, with model selection with ModelFinder and 1000 UFBoot2 approximations. For tests of selection, the ω ratio was estimated for each gene using the CODEML package of PAML, with individual gene trees being used. To detect amino acid sites under selection, four-site model comparisons were implemented (M0:M3, M1a:M2a, M7:M8, M8a:M8). These models have been extensively described elsewhere (Yang 2007). Briefly, the one ratio model (M0) allows for a single ω for all sites. The nearly neutral (M1a) has two categories of sites, ω = 1 and ω < 1. M2a (positive selection) has the same two categories as the M1a, with the addition of an ω > 1 category. The discrete (M3) model has three categories of sites for which ω can vary. M7 (‘beta’ neutral model) has eight site categories with eight ω taken from a discrete approximation of the beta distribution (range 0 – 1), meaning that the signal for positive selection cannot be detected. M8 (‘beta’ plus ω) has the same eight categories as the M7 model, plus and additional categories for which ω can vary from 0 to > 1. While the M8a (beta plus ωS=1) model is similar to the M8 model, with the ω_1_ being fixed at one. Multiple models and comparisons were used to test the robustness of the patterns observed. The M0:M3 tests whether ω varies between sites. The other comparisons test for positive selection, with the M1a:M2a being a more conservative test, the M7:M8 having more power and the M8a:M8 combining both power and robustness. Likelihood ratio tests (LRTs) between nested models, where ω is allowed to vary above one and the associated null model, allows inference of the selection pressures acting along the protein sequence (Yang 2007). P-values were generated from the LRT tests using the R function pchisq, with correction for multiple tests being carried out with the holm-bonferroni correction using the p.adjust function. Codons under positive selection were identified using the BEB (Bayes Empirical Bayes) method under the M2a model.

For visualisation of positively selected sites within the ANXA1, TNFSF4, CXCL16 and ANKRD17 protein structures, Phyre2 (Kelley et al. 2015) was used to predict protein models using a reference bat protein sequence (XP_036186722.1, XP_036199699.1, XP_036196999.1, XP_036191841.1). Where reliable models could be constructed, the resulting structure was used to predict relative solvent accessibility (RSA) with PolyView-2D (Porollo et al. 2004). Binding sites in the ANXA1 protein were identified by comparison against the Rat version of the protein (Uniprot Id: P07150). For transmembrane domain containing proteins, CXCL16 and TNFSF4, the location of transmembrane domains and signal peptides was carried out with Protter (Omasits et al. 2014).

### 6. Gene Ontology

Functional enrichment was tested using g:Profiler v. e109_eg56_p17_773ec798 (Raudvere et al. 2019). Enrichment tests for both overrepresentation and underrepresentation were performed using the list of 2,515 genes (Phylogenetic Dataset). Overrepresentation enrichment was analysed for the 38 genes identified as under selection in the Curated Gene Set. For all comparisons, human annotations were used with the background list of all annotated genes. GO enrichment was measured using a g:SCS multiple testing correction threshold of 0.05 and the GO:BP, KEGG, Reactome, and WikiPathways databases.

An overview of how the three datasets are related and which analyses were carried out are shown in Figure 1.

**Figure 1:**
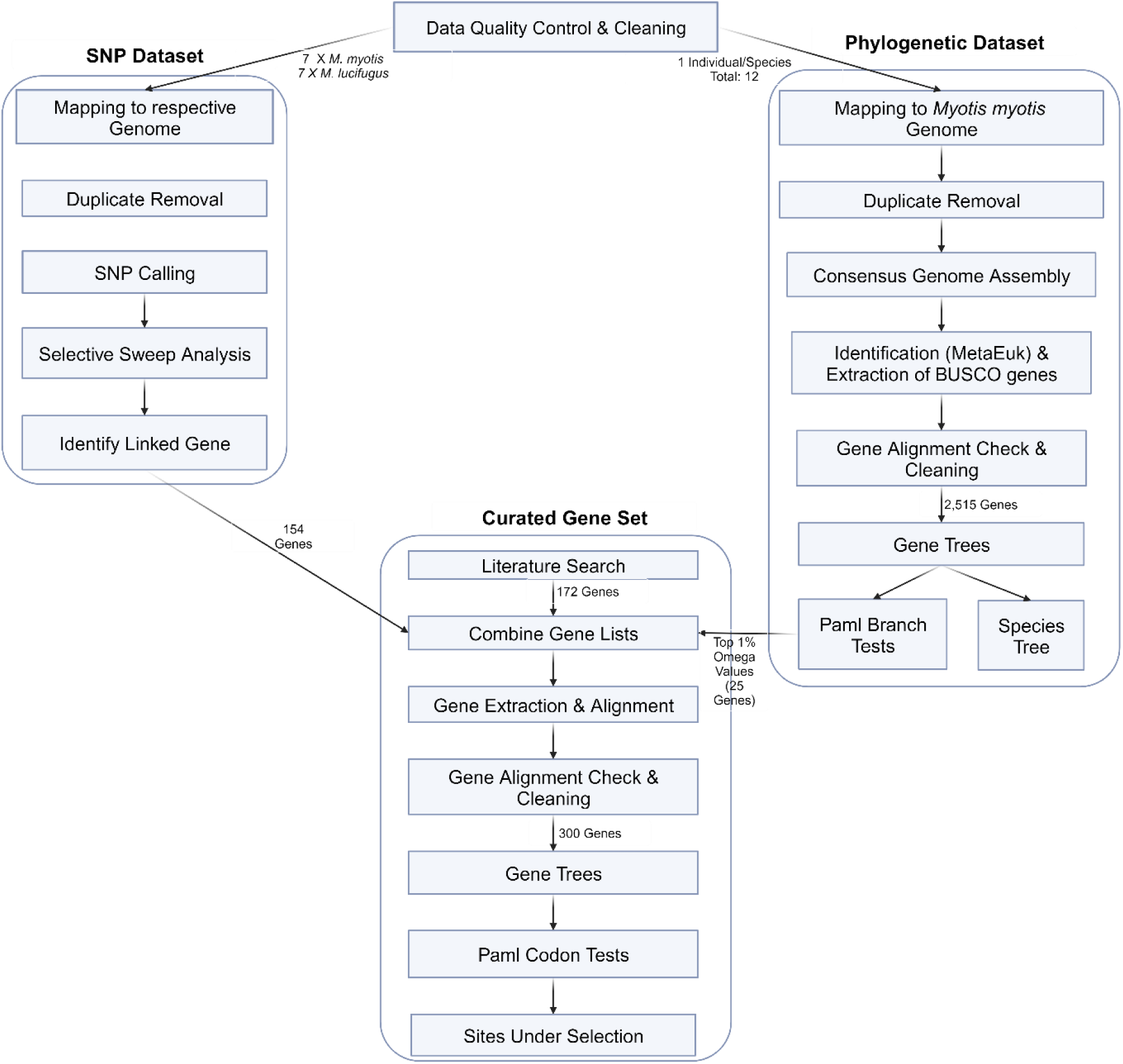
Overall workflow of the three main datasets and their associated analyses. Created with BioRender.com

## Results

We generated low coverage whole genome data for 18 *Myotis* specimens, encompassing 12 species. In addition, a further six *M. lucifugus* samples were downloaded from genbank (Supplementary Table 2). Mapping to the *M. myotis* reference genome resulted in an average mapping rate of 96% (Range: 85% – 99%) and an average genomic depth of coverage of 18X (Range: 6X – 53X). See Supplementary Table 6 for full sequencing statistics.

### Phylogenetic Dataset

Screening of single copy orthologues among our 12 *Myotis* species, resulted in a final dataset of 2,515 genes. Gene completeness ranged from 82 – 100%, with 2,713 of these genes being found in all 12 species (Supplementary Table 3). When this list of 2,515 genes was tested for functional enrichment (Supplementary Figure 1), we found that some significant enrichment was observed, but mostly for broad functional categories, such as GO:0050794 regulation of cellular process (padj = 2.632×10-17) and GO:0140053 mitochondrial gene expression (padj = 4.751×10-7). We also tested this gene list for underrepresentation and found that certain categories were significantly underrepresented, such as GO:0000244 spliceosomal tri-snRNP complex assembly (padj = 1.790×10-52) and GO:0007186 G protein-coupled receptor signalling pathway (padj = 2.978×10-4), but no immune-related categories were found (Supplementary Figure 2). We conclude that this list was not strongly biased in favour of any particular subset of functional pathways and that various immune pathways were sufficiently represented.

Species tree reconstruction (Figure 2) resulted in a topology with generally high support. Astral support values of 100% were estimated for all nodes, with the exception of the split between *M. daubentonii* and *M. bechsteinii*. Generally speaking, the relationships reconstructed agree with previous studies (Ruedi & Mayer 2001, Ruedi et al. 2013, Morales et al. 2019), with the primary difference in our topology being that *M. pequinus* is recovered as the sister species to *M. myotis* rather than *M. natteri*. We also see the mystacinus clade is hovering between the Eurasian and North American groups, a very different location to where it was assigned in the past (Ruedi et al. 2013) but has recently been subject to a shift in Morales et al. 2019.

**Figure 2:**
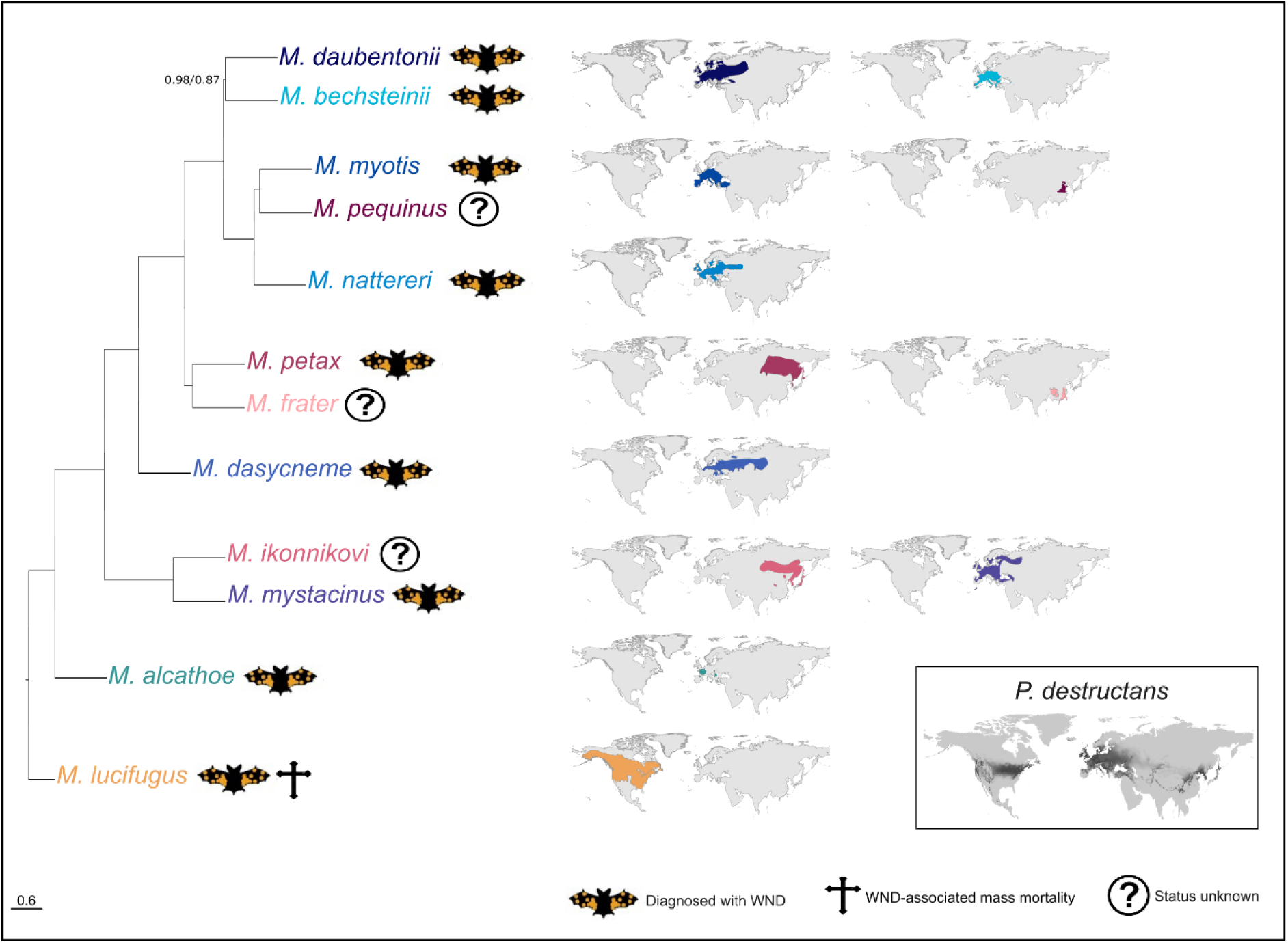
Phylogenetic relationships among 12 *Myotis* species based on species tree reconstruction using 2,515 genes. Support values represent Astral Support/bootstrap values from the two runs of ASTRAL-III, unless otherwise shown all nodes had 1.00/1.00 support. Distribution maps for each species are colour coded according to the taxa on the phylogeny, with the *P. destructans* distribution being shown in grey. The distribution maps are based on IUCN (International Union for Conservation of Nature, https://www.iucnredlist.org Accessed on 03 October 2023). Map of distribution range of *P. destructans* based on Blomberg et al. (2023).

To test for differences in selection in the branches leading to either *M. myotis* or *M. lucifugus,* independent branch tests were carried out. In each gene tree, the branch leading to either *M. myotis* or *M. lucifugus* was labelled as the foreground and allowed to have an independent ω ratio relative to the rest of the phylogeny. In all tests, no genes were found to have a statistically significant difference between the foreground and background ratios (Supplementary Table 7). A comparison of ω values under the neutral model for the phylogeny as a whole revealed strong patterns of purifying selection (ω < 1), with 18 genes having an ω value > 1 (average: 0.20, Range: 0.0001 – 2.12, Supplementary figure 3A). The highest gene with the highest ω value was *TNFSF4* (BUSCO code:188443at40674, ω of 2.12).

### SNP calling, selective sweeps detection and nucleotide diversity

With ANGSD we called after filtering 1,438,586 SNPs for *M. lucifugus* and 3,915,084 SNPs for *M. myotis*, and with GATK it was 4,304,169 and 11,757,539 SNPs, respectively (Supplementary Table 8). Of these, the final set of overlapping SNPs was for *Myotis lucifugus* 927,923 SNPs, and for *Myotis myoti*s 3,005,290 SNPs (Supplementary Table 8). In *Myotis lucifugus,* there were 464 outlier selective sweep windows which covered 17 genes (Supplementary Figure 4, Supplementary Table 9). In *Myotis myotis,* there were 1,499 outlier windows and 174 genes (Supplementary Figure 5, Supplementary Table 10). Three outlier genes were shared between the two species: *ARHGEF4, PLSCR1* and *PLOD2*. Nucleotide diversity, Theta Watterson, and Tajima’s D were higher in *Myotis myotis* than in *Myotis lucifugus* (Supplementary Figure 6A). Furthermore, some of the sweep regions matched with the low diversity regions in the whole genome level diversity plots (Supplementary Figure 6A and B).

### Codon test results for curated gene set

A final gene set of 300 protein coding genes were identified and extracted from our dataset. These 300 genes consisted of those genes identified in: I) Literature search, II) Selective Sweep analysis and III) overall ω values. The overlap between each of these categories is shown in Figure 3.

**Figure 3:**
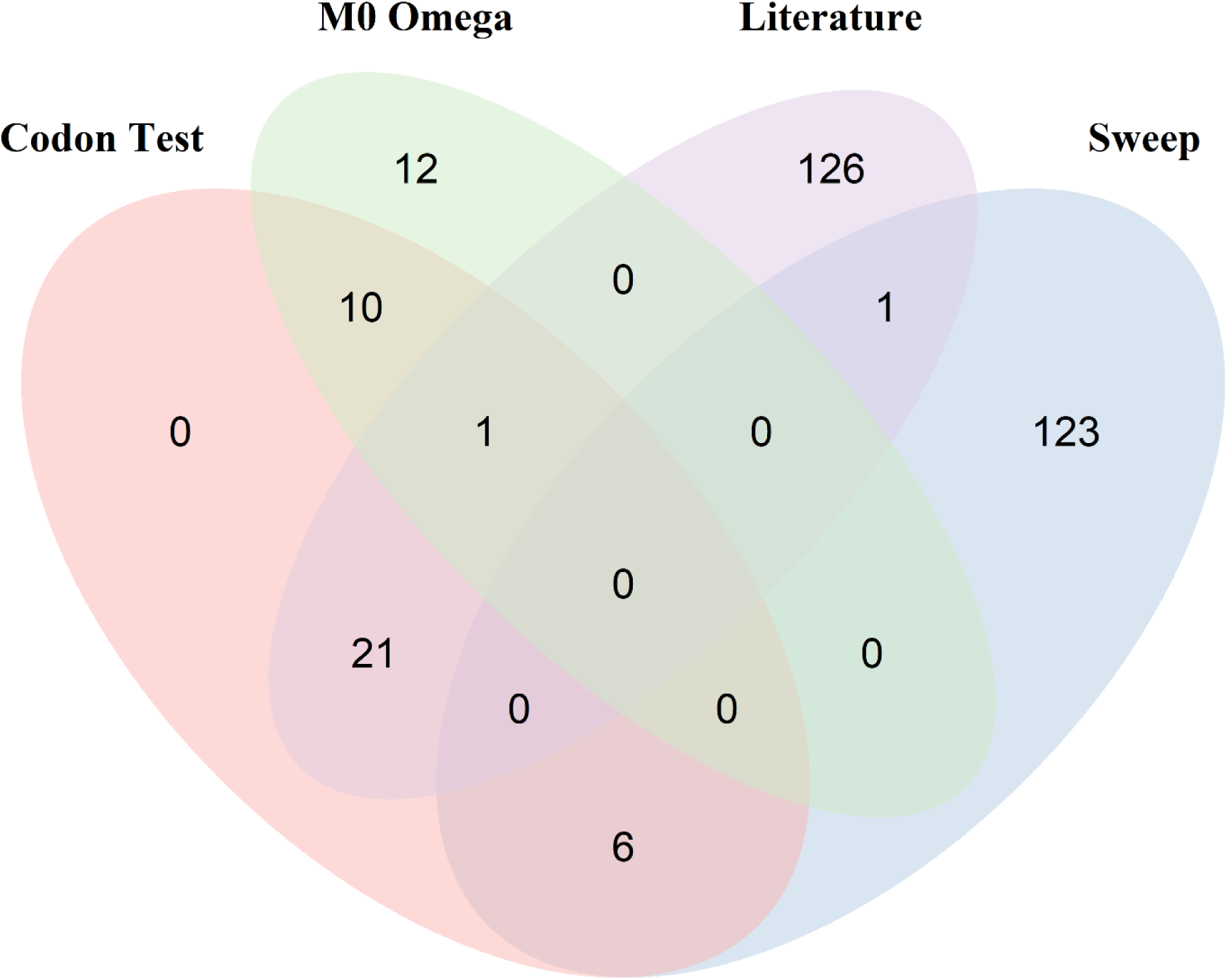
Venn Diagram showing the overlap between our various datasets and analyses. M0 Omega refers to the top 1% of omega values from the branch test null model; Sweep refers to the genes linked to the SNPs identified in our selective sweep analysis; Literature refers to all the genes identified from our literature search; and Codon Test refers to the 38 genes identified as under positive selection from our curated gene dataset.

Of the 300 genes, 270 were identified in all 12 species (Supplementary Table 5). To determine the selective pressures acting on these genes, an ω value was calculated for each orthologous gene set using site-based likelihood models. We found the majority of genes to be evolving under purifying selection when a single ω value is calculated across the entire coding region using the one-ratio (M0) model, mean = 0.32, range 0.0001 – 2.12 (Supplementary Figure 3B, Supplementary Table 11). Twenty-one genes have an ω > 1, indicating the protein as a whole is under positive selection (Supplementary Table 12A). As seen in the phylogenetic dataset, *TNFSF4* had the highest ω of 2.12. Rate heterogeneity among sites was tested using the M0:M3 comparison; significant among site variation was observed in 107 of the 300 genes tested. This, however, is not a test of positive selection. The three different model comparisons allow for the identification of positive selection among codons, each varying in their power and robustness. The genes identified in each comparison are given in Supplementary Table 12B. A total of 46 genes were identified in one or more comparisons, with 38 genes being identified in all three tests. In these 38 genes showing the strongest evidence for positive selection, we identified specific amino acids sites under significant selection. The number of sites identified as statistically significant under the robust M2a model ranges from 0 – 46 for each gene (Supplementary Table 13). Of the 38 genes showing the strongest signal for selection, six were from the selective sweep genes, 11 from the top 1% of omega values and 22 from the literature search (Figure 3).

The 38 genes that were identified with the strongest evidence for selection were further examined for functional enrichment using a statistical enrichment analysis and human gene ontology databases. We found that this gene list was highly enriched for genes with functional annotations related to many immune-related categories (Supplementary Table 14) and categories related to inflammatory immune responses were particularly common (Figure 4). For example, in the GO:Biological Process database, we found enrichment of GO:0045087 innate immune response (padj = 5.515×10-6), GO:0006954 inflammatory response (padj = 5.836×10-6), GO:0032637 interleukin-8 production (padj = 9.751×10-6) GO:0002250 adaptive immune response (padj = 3.393×10-4), and GO:0030097 hemopoiesis (padj = 3.735×10-3). In the KEGG database, we found enrichment for category KEGG:04613 Neutrophil extracellular trap formation (padj = 3.185×10-6); in the Reactome database, we found enrichment for categories involved in the TLR2/4 pathway, such as REAC:R-HSA-5602498 MyD88 deficiency (TLR2/4) (padj = 3.612×10-5); and in the WikiPathways database, we found enrichment of category WP:WP4493 Cells and molecules involved in local acute inflammatory response (padj = 2.953×10-5). Together, these functional enrichment results demonstrate that the majority of the genes identified under the strongest selection by our screens are related to inflammatory immune responses to the fungal pathogen mediated by both innate and adaptive mechanisms.

**Figure 4:**
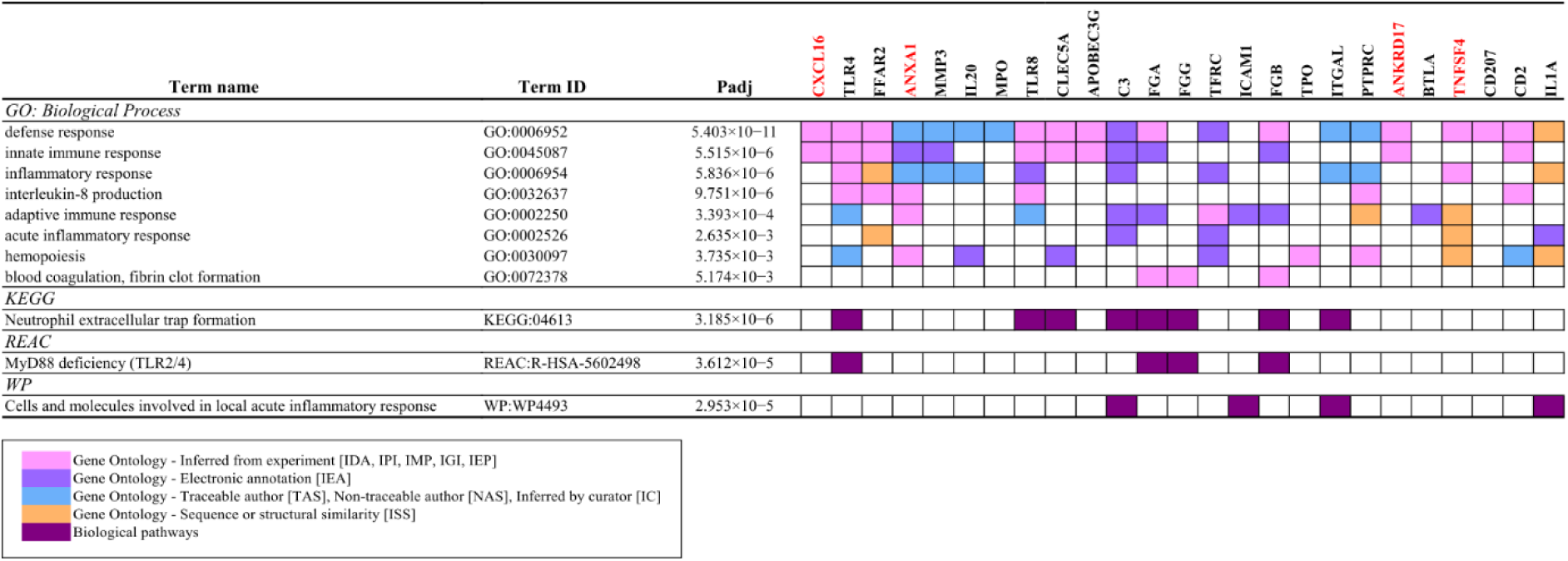
An overview of some of the key categories enriched in our 38 genes under positive selection. The four genes discussed in further detail are highlighted in red.

*TNFSF4* was the gene with the highest overall ω in the Phylogenetic and Curated Gene datasets, with an overall ω of 2.12. In addition, 9 sites were identified in our codon tests as being under positive selection, with ω values of ∼9 (Supplementary Table 13, Figure 5A). All nine sites are located in the extracellular portion of the protein (Supplementary Figure 7). Modelling of the three dimensional protein structure was successfully for ∼60% of the protein sequence (Supplementary Model 1, Figure 5B, a 360° view is shown in Supplementary File 1); assessing the solvent accessibility of the resulting model showed that residue 121 is fully exposed to the solvent, with an additional 3 sites (22, 64, 66) at least partially exposed, with RSA values > 5, and the remaining 5 being mostly buried within the protein structure (RSA < 5)(Supplementary Figure 8).

**Figure 5:**
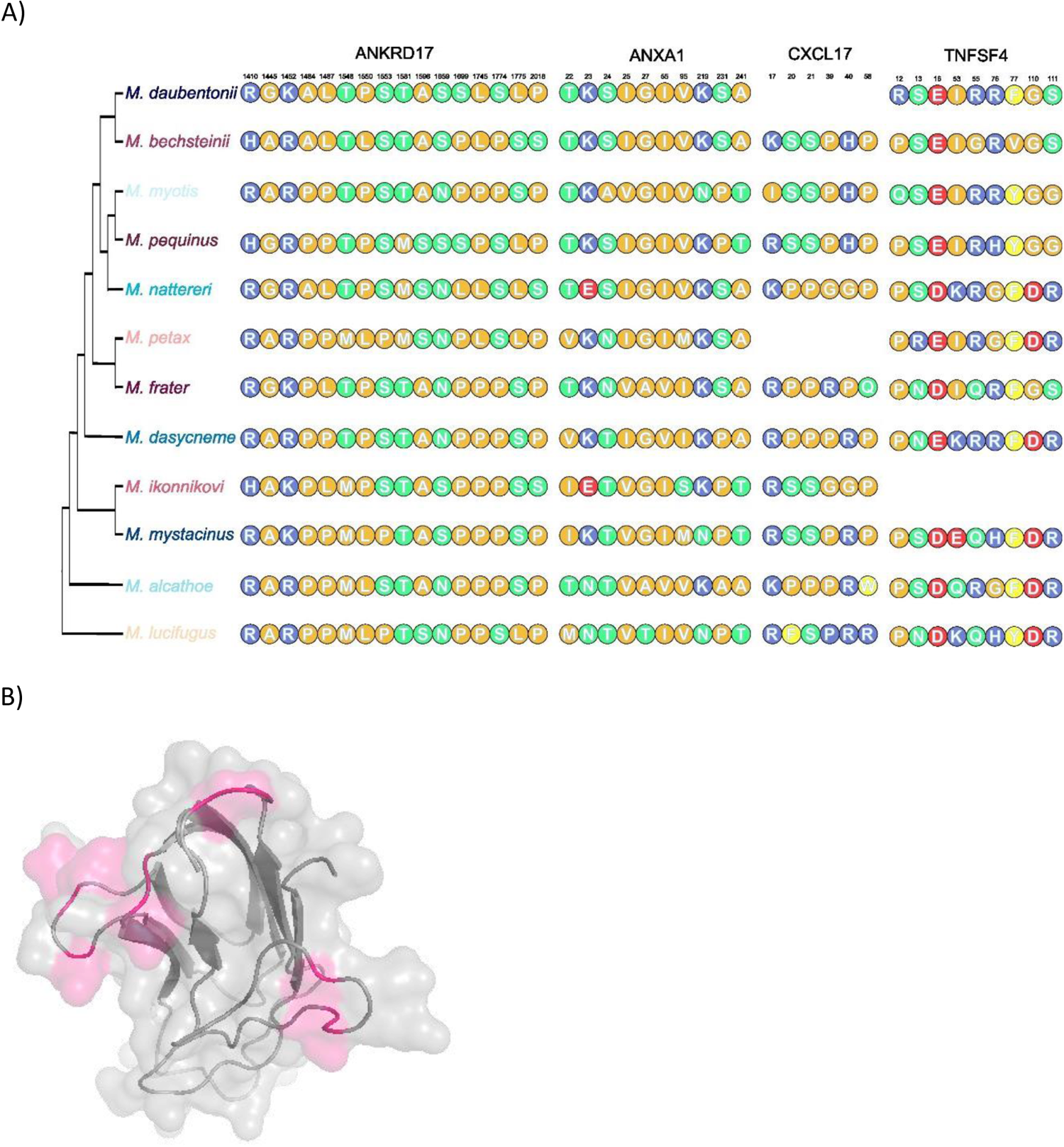
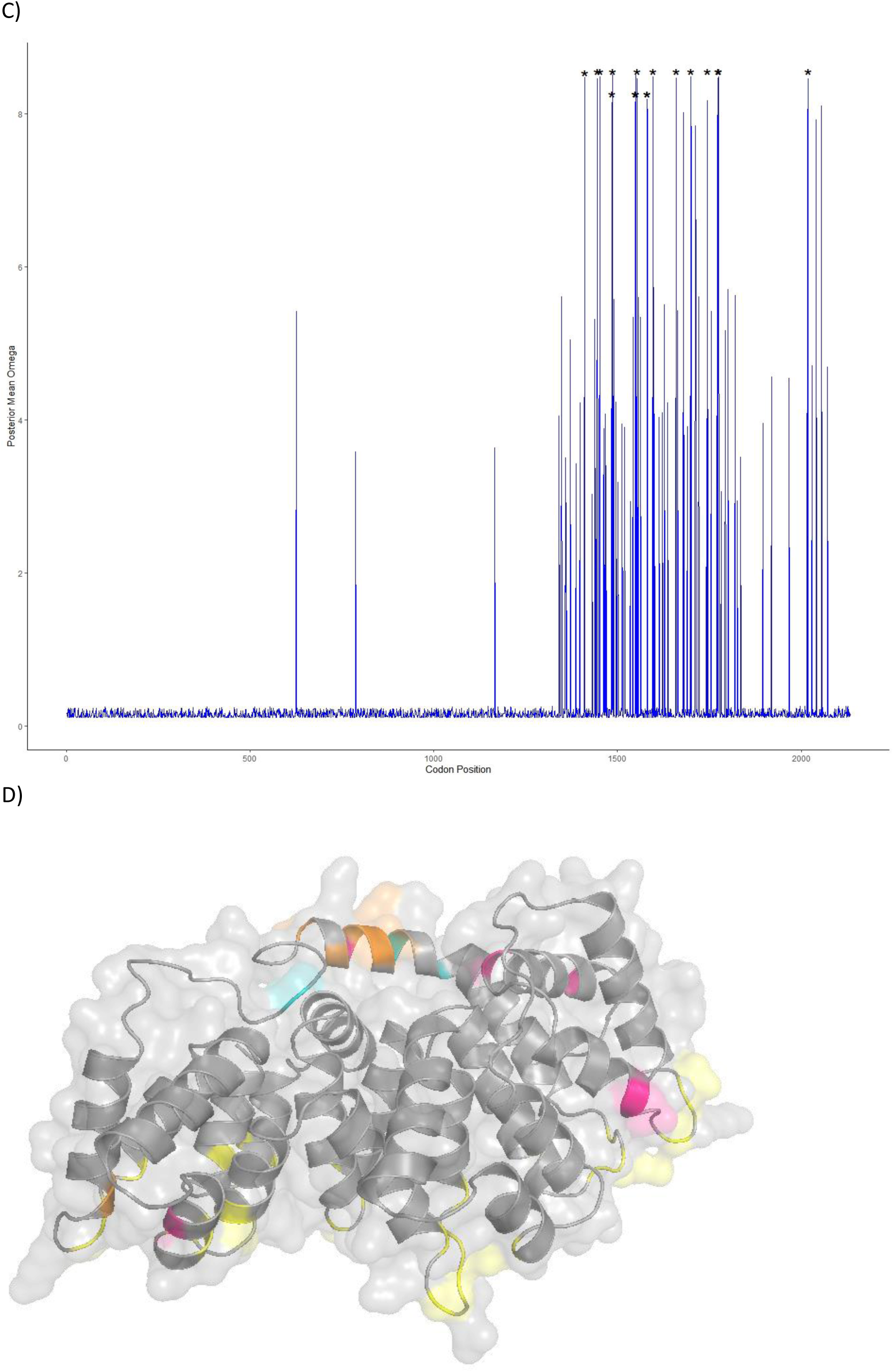
A) An overview of the sites under positive selection and their amino acid changes relative to their phylogenetic position for our four proteins (ANKRD17, ANXA1, CXCL16, TNFSF4) of interest. The phylogenetic tree shown is the same as that shown in Figure 2. B) Three-dimensional model of TNFSF4, with the residues under selection being highlighted in pink. C) Plot of omega value along the length of the ANKRD17 protein, positions with an * are those identified as being under positive selection. D) Three-dimensional model of ANXA1, with the calcium binding sites being shown in yellow, sites under positive selection have been coloured as follows: Pink: Codons under selection in our study; Orange: Codons under selection in both this study and (Harazim et al. 2018b); Aqua: Codons under selection in (Harazim et al. 2018b).

Among the 38 genes with strong evidence of positive selection, the gene *CXCL16* was the only gene to have originated from multiple datasets (being identified in the literature screen (Lilley et al. 2019) and among the genes with an ω > 1 in the Phylogenetic dataset). The protein as a whole has an ω of 1.4, with six codons being identified as being under positive selection (ω range: 4.6 – 4.8, Supplementary Table 13, Figure 5A). We were unable to model the structure of *CXCL16* with high confidence, however all six sites occur within the extracellular region of the protein (Supplementary Figure 9).

Of the 11 sweep genes also identified in the codon analysis, *ANKRD17* had the largest proportion of codons under positive selection. Overall, *ANKRD17* had an ω of 0.3. A total of 16 codons were identified as being under positive selection all with an ω value of ∼8 (Supplementary Table 13, Figure 5A). All of the 16 codons occur towards the end of the coding region (Figure 5C), with very little variation seen in approximately the first half of the coding sequence (Supplementary Figure 10). It was not possible to model the structure of this protein with high confidence, so no inference on where the 16 codons are located in the three-dimensional structure is possible.

Annexin A1 (*ANXA1)* was among the genes with positively selected sites, that had previously been identified in other studies (Harazim et al. 2018a, Lilley et al. 2019), with some evidence for codon specific selection occurring. Overall, *ANXA1* had an ω of 0.5, with 10 codons being identified as under positive selection with ω values of ∼7.8 (Supplementary Table 12 and 13, Figure 5A). Half of the sites identified are located in the N-terminal region of the protein (5 out of 10 sites identified are within the range 22 – 27). Relative solvent accessibility calculations based on the modelled structure (Supplementary Model 2, Figure 5D, a 360° view is shown in Supplementary File 2), show residue 27 is fully exposed to the solvent, while residues 22, 23 and 219 are partially exposed (RSA ≥ 5) and the remaining residues are either completely or mostly buried within the protein (Supplementary Figure 11).

## Discussion

Our comprehensive genomic study, utilising candidate genes sourced from a variety of data sets, detected strong positive selection in a suite of genes in Palearctic species of Myotis. Although not through an entirely unbiased approach, a considerable proportion of the genes have been associated with responses to fungal infection.

The species tree generated in this study using 2,515 genes generally agrees with the more recent studies on Myotis relationships. However, the tree produced for this study is limited due to the scope of the study and no novel results on evolutionary relationships can be inferred. The patterns of selection on the proteins the data set provide highlight overall functional constraint (ω < 1). This is a common occurrence with proteins generally being under purifying selection as a whole, with adaptation occurring within proteins at selected sites/regions (Nielsen & Yang 1998, Yang et al. 2000). Of the 2,515 genes tested, 21 had overall ω values indicative of positive selection. Of these, *TNFSF4* had the strongest signal for positive selection. We found no evidence for lineage specific selection associated with *M. myotis*, the species most associated with infection in the Palearctic. The same was evident for *M. lucifugus*, one of the most heavily affected species in North America, for which our individuals were sampled at the onset of the epizootic with no prior exposure.

We also looked at signals of selection within European and North American species, *Myotis myotis* and *Myotis lucifugus* respectively, using selective sweep analysis. A selective sweep is a process in which a beneficial mutation increases in frequency and becomes fixed in a population and this leads to a reduction in genetic diversity in the genomic region around the selected locus. Over a hundred genes were outliers in the *Myotis myotis* population, in contrast to the *Myotis lucifugus,* where only seventeen genes were in the sweep regions. Three genes were shared between the species, most notably phospholipid scramblase 1 (*PLSCR1*), an important gene in the antiviral responses (Xu et al. 2023). It is important to note that while selective sweep analyses can provide valuable insights into the genetic basis of adaptation, they have limitations, and the conclusions drawn should be interpreted with caution. In our case, sample size and sequencing coverage were the most important factors limiting the analyses. Other evolutionary processes and demographic factors as well can also influence patterns of genetic diversity, and it can be challenging to distinguish the causality between these. Therefore, multiple lines of evidence and additional studies are often necessary to confirm the effects of selection on specific loci.

We constructed a curated gene set of 300 proteins to investigate selection patterns within genes potentially associated with WNS tolerance. As with our phylogenetic dataset, most genes had an ω < 1, indicative of overall purifying selection. Twenty-one genes including *TNFSF4* showed signals for positive selection over the whole gene. Among our 300 genes, we found evidence for selection acting on at least one amino acid site in 38 genes.

We found that *TNFSF4* showed the most positive selection amongst the genes analysed in our study. Interestingly, this gene did not originate from the candidate set of genes gleaned from literature on WNS, but rather from the phylogenomic analysis that took an unbiased approach to identifying genes for our downstream analyses. This protein, tumor necrosis factor (ligand) superfamily, member 4, also known as OX-40L or CD252, is a potent activator of inflammatory signalling when it is expressed on antigen-presenting cells where it can activate OX-40 on T lymphocytes, NK cells, neutrophils, and others (Croft 2010). TNFR/TNF superfamily members can control diverse aspects of immune function. The mostly extracellular protein that is encoded by TNFSF4, in conjunction with its partner TNFSRF4, is associated with strong regulation of conventional CD4 and CD8 T cells, modulation of NKT cell and NK cell function, and mediation of cross-talk with professional antigen-presenting cells and diverse cell types such as mast cells, smooth muscle cells and endothelial cells. Additionally, TNFSF4-TNFSRF4 interactions alter the differentiation and activity of regulatory T cells (Croft 2010) and TNFSF4 mediates adhesion of activated T cells to endothelial cells during infection, inducing secretion of proinflammatory cytokines (Chiang et al. 2008). This gene clearly has a diverse function in the immune system but has also been specifically linked to responses to fungal infections. Several investigations have demonstrated that *TNFSF4* expression is upregulated in mice sensitized to *Aspergillus fumigatus* (Barrios et al. 2005) and that TNFSF4 polymorphisms are associated with the risk of developing invasive aspergillosis infection (Sánchez-Maldonado et al. 2021). In addition, blocking TNFSF4 has produced strong therapeutic effects in multiple animal models of autoimmune and inflammatory disease (Croft 2010)

*CXCL16* is another notable gene, filtering through to the final dataset from both the top 1% of genes in the phylogenetic dataset, and the literature search, being previously observed as an upregulated gene in naïve infected *M. lucifugus* (Lilley et al. 2019). The protein, expressed as either a soluble or a transmembrane form, is a marker of inflammation in humans and is produced by monocytes/macrophages, B cells, dendritic cells, keratinocytes, and endothelial cells (Hase et al. 2006). As the sole ligand for the receptor CXCR6, soluble CXCL16 promotes the directional migration of CXCR6^+^ cells, such as CD4^+^ effector memory T cells and natural killer T-cells (Wilbanks et al. 2001), with expression being induced by inflammatory cytokines (Abel et al. 2004). All sites under selection in our study are in the extracellular section of the transmembrane polypeptide, indicating possible functional changes in the interactions between CXCL16 and its receptor, CXCR6.

ANKRD17 is an ankyrin repeat protein, which has previously been reported to be an important regulator of the cell cycle (Deng et al. 2009) and may also be involved in innate immune activation by viruses and bacteria (Wang et al. 2012, Menning & Kufer 2013). The codons under selection for this gene are clustered in the C-terminus of the protein, outside of the region of ankyrin repeats that has been shown to bind to the NLR-family pattern-recognition receptor, Nod2 (Menning & Kufer 2013), and in the region that is putatively responsible for activation of RIG-I-like receptors (Wang et al. 2012). This may indicate some cross-talk between antifungal and antiviral innate signalling pathways, or it could be related to the presence of viral coinfections in some bats with WNS (Davy et al. 2018).

Our literature search identified several studies that implicated ANXA1, Annexin A1, a protein that is a well-characterized anti-inflammatory inhibitor of cytosolic phospholipase A2 (Gerke & Moss 2002). In addition, ANXA1 is a known target for positive selection in *Myotis* infected by *P. destructans* (Harazim et al. 2018a), and shows increased transcription in hibernating, infected Nearctic *M. lucifugus* compared to uninfected conspecifics, or Palearctic *M. myotis* (Lilley et al. 2019). Harazim et al. 2018 suggested ANXA1 may act via two different routes in bats infected by *P. destructans*. First, ANXA1 has the potential to down-regulate the immune responses initiated by bats arousing from torpor, which can lead to immunopathology if left unchecked. ANXA1 appears to regulate the neutrophil response under fungal infection conditions, altering lipid membrane organization and metabolism (Sanches et al. 2021). ANXA1 has also been found to participate in adaptive immunity against chronic infectious disease (Vanessa et al. 2015), by directing the immune response towards a Th1/Th17 response, a response that is associated not only with clearing fungal infection in normothermic mammals, but also the mortality evidenced in susceptible Nearctic populations of *M. lucifugus* infected by *P. destructans* (Lilley et al. 2017). Second, ANXA1 also appears to play a role in wound repair and epithelial recovery, extending its importance beyond the acute phase of inflammation to the equally important healing phase (Leoni et al. 2015, Fuller et al. 2020). Our study found five of the seven sites under positive selection in the study by Harazim et al. (2019), with an addition of four previously undescribed sites. All previously identified sites were confined to the N-terminal region of the protein, where ANXA1 is known to bind to S100A11 and inhibit phospholipase A2 (Mailliard et al. 1996, Sakaguchi et al. 2007). We found additional evidence for selection at three sites towards the C-terminal end of the protein, whose function is not as well characterized.

Our combined approach to identify genes involved in adaptation to WNS revealed a preponderance of genes with known functions involved in defence responses, including both innate and adaptive immune responses. In addition to the specific genes described above, we found strong evidence that pathways that activate inflammatory responses and neutrophil recruitment are under selection. This is highly consistent with the observed pathology associated with WNS in naive populations, both during hibernation and after emergence (Meteyer et al. 2012). IL-8 signalling appears to be among the inflammatory pathways that are particularly important, and this cytokine is known to recruit neutrophils to sites of infection and enhance their activation, a process that likely generates significant immunopathology. Other pathways that appear to be involved in adaptation included coagulation and stress responses, although it is not clear how increased or decreased activation of these pathways would enhance disease tolerance. Presumably the driving force of natural selection is to suppress the inflammatory responses that lead to immunopathology, or at least delay them until it is more energetically favourable to mount a resistance response.

## Conclusions

A volume of research over the last decade indicates not only the significance of immunopathology in the cascade of physiological events that build up to mortality associated with WNS (Field et al. 2015, 2018, Lilley et al. 2017), but also the lack of immunopathology in infected hosts in the Palearctic (Fritze et al. 2019, 2021, Lilley et al. 2019, Hecht-Höger et al. 2020). This is suggestive of the evolution of tolerance in Palearctic bats through coevolution over an extended period of time (Leopardi et al. 2015, Whiting-Fawcett et al. 2021). Selection towards tolerance to the fungal infection in the Palearctic has been suggested (Whiting-Fawcett et al. 2021) and indicated at a genomic level (Harazim et al. 2018a), but here, through an exhaustive approach, we can state with higher confidence that genes and pathways associated with fungal infections, and particularly those involved in infection by *P. destructans*, have been under positive selection in Palearctic species of Myotis. Further research through e.g., proteomics and immunological modelling is needed to elucidate exactly how the observed selection affects response to infection. Nevertheless, identifying genes, and their variants will allow for the estimation of survival in North American affected species through population genetic approaches to assist in implementing conservation measures.

## Acknowledgements

The authors wish to acknowledge CSC-IT Center for Science, Finland, for computational resources.

## Data Accessibility

All raw data generated in this study are accessible at # under #. All final gene alignments, and gene trees can be found on Zenodo under the doi: XXXX. All supplementary files are available at: https://www.dropbox.com/scl/fo/nsf73e547racaepd2tv4m/h?rlkey=csd5m7d17yv8qi0dtd92bp533&dl=0

## References

1. Abel S, Hundhausen C, Mentlein R, Schulte A, Berkhout TA, Broadway N et al. (2004) The Transmembrane CXC-Chemokine Ligand 16 Is Induced by IFN-γ and TNF-α and Shed by the Activity of the Disintegrin-Like Metalloproteinase ADAM10 1. The Journal of Immunology 172: 6362–6372.

2. Alachiotis N, Pavlidis P (2018) RAiSD detects positive selection based on multiple signatures of a selective sweep and SNP vectors. Communications Biology 1: 1–11.

3. Andrews S (2015) FastQC A Quality Control tool for High Throughput Sequence Data.

4. Ayres JS, Schneider DS (2012) Tolerance of infections. Annual Review of Immunology 30: 271– 294.

5. Baker RE, Mahmud AS, Miller IF, Rajeev M, Rasambainarivo F, Rice BL et al. (2022) Infectious disease in an era of global change. Nature Reviews Microbiology 20: 193–205.

6. Barrios CS, Johnson BD, D. Henderson J, Fink JN, Kelly KJ, Kurup VP (2005) The costimulatory molecules CD80, CD86 and OX40L are up-regulated in Aspergillus fumigatus sensitized mice. Clinical and Experimental Immunology 142: 242–250.

7. Baylis M, Risley C (2012) Infectious DiseasesInfectious Diseases, Climate Change Effectsinfectious diseaseclimate change Effectson. In: Meyers RA (ed) Encyclopedia of Sustainability Science and Technology, 5358–5378. Springer, New York, NY.

8. Blehert DS, Hicks AC, Behr M, Meteyer CU, Berlowski-Zier BM, Buckles EL et al. (2009) Bat white-nose syndrome: an emerging fungal pathogen? Science 323: 227–227.

9. Blomberg AS, Lilley TM, Fritze M, Puechmaille SJ (2023) Climatic factors and host species composition at hibernation sites drive the incidence of bat fungal disease. : 2023.02.27.529820.

10. Bolger AM, Lohse M, Usadel B (2014) Trimmomatic: A flexible trimmer for Illumina Sequence Data.

11. Castresana J (2000) Selection of Conserved Blocks from Multiple Alignments for Their Use in Phylogenetic Analysis. Molecular Biology and Evolution 17: 540–552.

12. Cheng TL, Gerson A, Moore MS, Reichard JD, DeSimone J, Willis CKR, Frick WF, Kilpatrick AM (2019) Higher fat stores contribute to persistence of little brown bat populations with white-nose syndrome. Journal of Animal Ecology 88: 561–600.

13. Chiang LY, Sheppard DC, Gravelat FN, Patterson TF, Filler SG (2008) Aspergillus fumigatus Stimulates Leukocyte Adhesion Molecules and Cytokine Production by Endothelial Cells In Vitro and during Invasive Pulmonary Disease. Infection and Immunity 76: 3429–3438.

14. Croft M (2010) Control of Immunity by the TNFR-Related Molecule OX40 (CD134). Annual Review of Immunology 28: 57–78.

15. Cunningham AA, Daszak P, Wood JLN (2017) One Health, emerging infectious diseases and wildlife: two decades of progress? Philosophical Transactions of the Royal Society B: Biological Sciences 372: 20160167.

16. Dainat J, Murray K, Hereñú D, Davis E, Crouch K, Sol L et al. (2023) AGAT: Another Gff Analysis Toolkit to handle annotations in any GTF/GFF format. (Version v0.9.2).

17. Danecek P, Auton A, Abecasis G, Albers CA, Banks E, DePristo MA et al. (2011) The variant call format and VCFtools. Bioinformatics 27: 2156–2158.

18. Danecek P, Bonfield JK, Liddle J, Marshall J, Ohan V, Pollard MO et al. (2021) Twelve years of SAMtools and BCFtools. GigaScience 10: giab008.

19. Daszak P, Cunningham AA, Hyatt AD (2000) Emerging infectious diseases of wildlife-threats to biodiversity and human health. Science 287: 443–449.

20. Davy CM, Donaldson ME, Subudhi S, Rapin N, Warnecke L, Turner JM et al. (2018) White-nose syndrome is associated with increased replication of a naturally persisting coronaviruses in bats. Scientific Reports 8: 15508.

21. Deng M, Li F, Ballif BA, Li S, Chen X, Guo L, Ye X (2009) Identification and Functional Analysis of a Novel Cyclin E/Cdk2 Substrate Ankrd17 *. Journal of Biological Chemistry 284: 7875–7888.

22. Field KA, Johnson JS, Lilley TM, Reeder SM, Rogers EJ, Behr MJ, Reeder DM (2015) The white-nose syndrome transcriptome: activation of anti-fungal host responses in wing tissue of hibernating bats. PLoS Pathogens 11: 1–29.

23. Field KA, Sewall BJ, Prokkola JM, Turner GG, Gagnon MF, Lilley TM et al. (2018) Effect of torpor on host transcriptomic responses to a fungal pathogen in hibernating bats. Molecular Ecology 27: 3727–3743.

24. Fischer NM, Dool SE, Puechmaille SJ (2020) Seasonal patterns of *Pseudogymnoascus destructans* germination indicate host – pathogen coevolution. Biology Letters 16: 1–5.

25. Flieger M, Bandouchova H, Cerny J, Chudíčková M, Kolarik M, Kovacova V et al. (2016) Vitamin B2 as a virulence factor in *Pseudogymnoascus destructans* skin infection. Scientific Reports 6: 33200.

26. Frick WF, Johnson E, Cheng TL, Lankton JS, Warne R, Dallas J et al. (2022) Experimental inoculation trial to determine the effects of temperature and humidity on White-nose Syndrome in hibernating bats. Scientific Reports 12: 971.

27. Frick WF, Puechmaille SJ, Hoyt JR, Nickel BA, Langwig KE, Foster JT et al. (2015) Disease alters macroecological patterns of North American bats. Global Ecology and Biogeography 24: 741– 749.

28. Frick WF, Puechmaille SJ, Willis CKR (2016) White-nose syndrome in bats. In: Voigt CC, Kingston T (eds) Bats in the Anthropocene: Conservation of bats in a changing world, 245–262. Springer International Publishing, Cham.

29. Fritze M, Costantini D, Fickel J, Wehner D, Czirják GÁ, Voigt CC (2019) Immune response of hibernating European bats to a fungal challenge. Biology Open 8: 1–10.

30. Fritze M, Puechmaille SJ, Costantini D, Fickel J, Voigt CC, Czirják GÁ (2021) Determinants of defence strategies of a hibernating European bat species towards the fungal pathogen *Pseudogymnoascus destructans*. Developmental & Comparative Immunology: 104017.

31. Fuller NW, McGuire LP, Pannkuk EL, Blute T, Haase CG, Mayberry HW, Risch TS, Willis CKR (2020) Disease recovery in bats affected by white-nose syndrome. Journal of Experimental Biology 223: 1–12.

32. Gerke V, Moss SE (2002) Annexins: From Structure to Function. Physiological Reviews 82: 331– 371.

33. Harazim M, Horáček I, Jakešová L, Luermann K, Moravec JC, Morgan S et al. (2018a) Natural selection in bats with historical exposure to white-nose syndrome. BMC Zoology 3: 1–13.

34. Harazim M, Horacek I, Jakesova L, Luermann K, Moravec J, Morgan S et al. (2018b) Natural selection in bats with historical exposure to white-nose syndrome. BMC ZOOLOGY 3.

35. Hase K, Murakami T, Takatsu H, Shimaoka T, Iimura M, Hamura K et al. (2006) The Membrane-Bound Chemokine CXCL16 Expressed on Follicle-Associated Epithelium and M Cells Mediates Lympho-Epithelial Interaction in GALT1. The Journal of Immunology 176: 43–51.

36. Hecht-Höger AM, Beate CB, Krause E, Meschede A, Krahe R, Voigt CC, Greenwood AD, Czirják GÁ (2020) Plasma proteomic profiles differ between European and North American myotid bats colonized by *Pseudogymnoascus destructans*. Molecular Ecology 29: 1745–1755.

37. Hoang DT, Chernomor O, von Haeseler A, Minh BQ, Vinh LS (2018) UFBoot2: Improving the Ultrafast Bootstrap Approximation. Molecular Biology and Evolution 35: 518–522.

38. Hoyt JR, Sun K, Parise KL, Lu G, Langwig KE, Jiang T et al. (2015) Widespread Bat White-Nose Syndrome Fungus, Northeastern China. Emerging Infectious Diseases 22.

39. Joshi N, Fass J (2011) Sickle: A sliding-window, adaptive, quality-based trimming tool for FastQ files (Version 1.33) [Software]. Available at https://github.com/najoshi/sickle.

40. Kalyaanamoorthy S, Minh BQ, Wong TKF, von Haeseler A, Jermiin LS (2017) ModelFinder: fast model selection for accurate phylogenetic estimates. Nature Methods 14: 587–589.

41. Karlsson EK, Kwiatkowski DP, Sabeti PC (2014) Natural selection and infectious disease in human populations. Nature Reviews. Genetics 15: 379–393.

42. Katoh K, Standley DM (2013) MAFFT Multiple Sequence Alignment Software Version 7: Improvements in Performance and Usability. Molecular Biology and Evolution 30: 772–780.

43. Kelley LA, Mezulis S, Yates CM, Wass MN, Sternberg MJE (2015) The Phyre2 web portal for protein modeling, prediction and analysis. Nature Protocols 10: 845–858.

44. Kim SY, Lohmueller KE, Albrechtsen A, Li Y, Korneliussen T, Tian G et al. (2011) Estimation of allele frequency and association mapping using next-generation sequencing data. BMC Bioinformatics 12: 231.

45. Korneliussen TS, Albrechtsen A, Nielsen R (2014) ANGSD: Analysis of Next Generation Sequencing Data. BMC Bioinformatics 15: 356.

46. Kriventseva EV, Kuznetsov D, Tegenfeldt F, Manni M, Dias R, Simão FA, Zdobnov EM (2019) OrthoDB v10: sampling the diversity of animal, plant, fungal, protist, bacterial and viral genomes for evolutionary and functional annotations of orthologs. Nucleic Acids Research 47: D807–D811.

47. Langmead B, Salzberg SL (2012) Fast gapped-read alignment with Bowtie 2. Nature Methods 9: 357–359.

48. Leoni G, Neumann P-A, Kamaly N, Quiros M, Nishio H, Jones HR et al. (2015) Annexin A1-containing extracellular vesicles and polymeric nanoparticles promote epithelial wound repair. The Journal of Clinical Investigation 125: 1215–1227.

49. Leopardi S, Blake D, Puechmaille SJ (2015) White-nose syndrome fungus introduced from Europe to North America. Current Biology 25: R217–R219.

50. Levy Karin E, Mirdita M, Söding J (2020) MetaEuk—sensitive, high-throughput gene discovery, and annotation for large-scale eukaryotic metagenomics. Microbiome 8: 48.

51. Lilley TM, Prokkola JM, Blomberg AS, Paterson S, Johnson JS, Turner GG et al. (2019) Resistance is futile: RNA-sequencing reveals differing responses to bat fungal pathogen in Nearctic *Myotis lucifugus* and Palearctic *Myotis myotis*. Oecologia: 295–309.

52. Lilley TM, Prokkola JM, Johnson JS, Rogers EJ, Gronsky S, Kurta A, Reeder DM, Field KA (2017) Immune responses in hibernating little brown myotis (*Myotis lucifugus*) with white-nose syndrome. Proc. R. Soc. B 284: 20162232.

53. Lilley TM, Wilson IW, Field KA, Reeder DM, Vodzak ME, Turner GG et al. (2020) Genome-Wide Changes in Genetic Diversity in a Population of Myotis lucifugus Affected by White-Nose Syndrome. G3: *Genes, Genomes, Genetics* 10: 2007–2020.

54. Mailliard WS, Haigler HT, Schlaepfer DD (1996) Calcium-dependent binding of S100C to the N-terminal domain of annexin I. The Journal of Biological Chemistry 271: 719–725.

55. Manni M, Berkeley MR, Seppey M, Simão FA, Zdobnov EM (2021) BUSCO Update: Novel and Streamlined Workflows along with Broader and Deeper Phylogenetic Coverage for Scoring of Eukaryotic, Prokaryotic, and Viral Genomes. Molecular Biology and Evolution 38: 4647–4654.

56. Martin M (2011) Cutadapt removes adapter sequences from high-throughput sequencing reads. EMBnet.journal 17: 10–12.

57. Martinkova N, Backor P, Bartonicka T, Blazkova P, Cerveny J, Falteisek L et al. (2010) Increasing incidence of *Geomyces* destructans fungus in bats from the Czech Republic and Slovakia. Plos One 5: e13853.

58. McGuire LP, Mayberry HW, Willis CKR (2017) White-nose syndrome increases torpid metabolic rate and evaporative water loss in hibernating bats. *American Journal of Physiology-Regulatory*, Integrative and Comparative Physiology 313: R680–R686.

59. Menning M, Kufer TA (2013) A role for the Ankyrin repeat containing protein Ankrd17 in Nod1- and Nod2-mediated inflammatory responses. FEBS letters 587: 2137–2142.

60. Meteyer CU, Barber D, Mandl JN (2012) Pathology in euthermic bats with white nose syndrome suggests a natural manifestation of immune reconstitution inflammatory syndrome. Virulence 3.

61. Meteyer CU, Buckles EL, Blehert DS, Hicks AC, Green DE, Shearn-Bochsler V, Thomas NJ, Gargas A, Behr MJ (2009) Histopathologic criteria to confirm white-nose syndrome in bats. Journal of Veterinary Diagnostic Investigation 21: 411–414.

62. Minh BQ, Schmidt HA, Chernomor O, Schrempf D, Woodhams MD, von Haeseler A, Lanfear R (2020) IQ-TREE 2: New Models and Efficient Methods for Phylogenetic Inference in the Genomic Era. Molecular Biology and Evolution 37: 1530–1534.

63. Morales AE, Ruedi M, Field K, Carstens BC (2019) Diversification rates have no effect on the convergent evolution of foraging strategies in the most speciose genus of bats, Myotis*. Evolution 73: 2263–2280.

64. Nielsen R, Yang Z (1998) Likelihood Models for Detecting Positively Selected Amino Acid Sites and Applications to the HIV-1 Envelope Gene. Genetics 148: 929–936.

65. Omasits U, Ahrens CH, Müller S, Wollscheid B (2014) Protter: interactive protein feature visualization and integration with experimental proteomic data. Bioinformatics 30: 884–886.

66. Picard Tools – By Broad Institute

67. Porollo AA, Adamczak R, Meller J (2004) POLYVIEW: a flexible visualization tool for structural and functional annotations of proteins. Bioinformatics 20: 2460–2462.

68. Puechmaille SJ, Wibbelt G, Korn V, Fuller H, Forget F, Mühldorfer K et al. (2011) Pan-European distribution of white-nose syndrome fungus (*Geomyces destructans*) not associated with mass mortality. PLoS ONE 6: e19167.

69. Quinlan AR, Hall IM (2010) BEDTools: a flexible suite of utilities for comparing genomic features. Bioinformatics 26: 841–842.

70. R Core Team (2022) R: A language and environment for statistical computing. R Foundation for Statistical Computing, Vienna, Austria. http://www.R-project.org/.

71. Raudvere U, Kolberg L, Kuzmin I, Arak T, Adler P, Peterson H, Vilo J (2019) g:Profiler: a web server for functional enrichment analysis and conversions of gene lists (2019 update). Nucleic Acids Research 47: W191–W198.

72. Reeder DM, Frank CL, Turner GG, Meteyer CU, Kurta A, Britzke ER et al. (2012) Frequent arousal from hibernation linked to severity of infection and mortality in bats with white-nose syndrome. PLoS ONE 7.

73. Ruedi M, Mayer F (2001) Molecular Systematics of Bats of the Genus Myotis (Vespertilionidae) Suggests Deterministic Ecomorphological Convergences. Molecular Phylogenetics and Evolution 21: 436–448.

74. Ruedi M, Stadelmann B, Gager Y, Douzery EJP, Francis CM, Lin L-K, Guillén-Servent A, Cibois A (2013) Molecular phylogenetic reconstructions identify East Asia as the cradle for the evolution of the cosmopolitan genus Myotis (Mammalia, Chiroptera). Molecular Phylogenetics and Evolution 69: 437–449.

75. Sakaguchi M, Murata H, Sonegawa H, Sakaguchi Y, Futami J, Kitazoe M, Yamada H, Huh N (2007) Truncation of Annexin A1 Is a Regulatory Lever for Linking Epidermal Growth Factor Signaling with Cytosolic Phospholipase A2 in Normal and Malignant Squamous Epithelial Cells *. Journal of Biological Chemistry 282: 35679–35686.

76. Sanches JM, Rossato L, Lice I, Alves de Piloto Fernandes AM, Bueno Duarte GH, Rosini Silva AA, de Melo Porcari A, de Oliveira Carvalho P, Gil CD (2021) The role of annexin A1 in Candida albicans and Candida auris infections in murine neutrophils. Microbial Pathogenesis 150: 104689.

77. Sánchez-Maldonado JM, Moñiz-Díez A, ter Horst R, Campa D, Cabrera-Serrano AJ, Martínez-Bueno M et al. (2021) Polymorphisms within the TNFSF4 and MAPKAPK2 Loci Influence the Risk of Developing Invasive Aspergillosis: A Two-Stage Case Control Study in the Context of the aspBIOmics Consortium. Journal of Fungi 7: 4.

78. Schmieder R, Edwards R (2011) Quality control and preprocessing of metagenomic datasets. Bioinformatics 27: 863–864.

79. Shumate A, Salzberg SL (2021) Liftoff: accurate mapping of gene annotations. Bioinformatics 37: 1639–1643.

80. Stelzer G, Rosen N, Plaschkes I, Zimmerman S, Twik M, Fishilevich S et al. (2016) The GeneCards Suite: From Gene Data Mining to Disease Genome Sequence Analyses. Current Protocols in Bioinformatics 54: 1.30.1-1.30.33.

81. Suyama M, Torrents D, Bork P (2006) PAL2NAL: robust conversion of protein sequence alignments into the corresponding codon alignments. Nucleic Acids Research 34: W609– W612.

82. Vanessa KHQ, Julia MG, Wenwei L, Michelle ALT, Zarina ZRS, Lina LHK, Sylvie A (2015) Absence of Annexin A1 impairs host adaptive immunity against Mycobacterium tuberculosis in vivo. Immunobiology 220: 614–623.

83. Wang Y, Tong X, Li G, Li J, Deng M, Ye X (2012) Ankrd17 positively regulates RIG-I-like receptor (RLR)-mediated immune signaling. European Journal of Immunology 42: 1304–1315.

84. Warnecke L, Turner JM, Bollinger TK, Lorch JM, Misra V, Cryan PM, Wibbelt G, Blehert DS, Willis CKR (2012) Inoculation of bats with European *Geomyces destructans* supports the novel pathogen hypothesis for the origin of white-nose syndrome. Proceedings of the National Academy of Sciences of the United States of America 109: 6999–7003.

85. Whiting-Fawcett F, Field KA, Puechmaille SJ, Blomberg AS, Lilley TM (2021) Heterothermy and antifungal responses in bats. Current Opinion in Microbiology 62: 61–67.

86. Wilbanks A, Zondlo SC, Murphy K, Mak S, Soler D, Langdon P, Andrew DP, Wu L, Briskin M (2001) Expression cloning of the STRL33/BONZO/TYMSTRligand reveals elements of CC, CXC, and CX3C chemokines. Journal of Immunology (Baltimore, Md.: 1950) 166: 5145–5154.

87. Wilson DE, Mittermeier RA (eds) (2019) Handbook of the Mammals of the World. Vol. 9. Bats. Lynx Edicions, Barcelona.

88. Xu D, Jiang W, Wu L, Gaudet RG, Park E-S, Su M et al. (2023) PLSCR1 is a cell-autonomous defence factor against SARS-CoV-2 infection. Nature 619: 819–827.

89. Yang Z (2007) PAML 4: Phylogenetic Analysis by Maximum Likelihood. Molecular Biology and Evolution 24: 1586–1591.

90. Yang Z, Bielawski JP (2000) Statistical methods for detecting molecular adaptation. Trends in Ecology & Evolution 15: 496–503.

91. Yang Z, Nielsen R, Goldman N, Pedersen AM (2000) Codon-substitution models for heterogeneous selection pressure at amino acid sites. Genetics 155: 431–449.

92. Zhang C, Rabiee M, Sayyari E, Mirarab S (2018) ASTRAL-III: polynomial time species tree reconstruction from partially resolved gene trees. BMC Bioinformatics 19: 153.

93. Zukal J, Bandouchova H, Brichta J, Cmokova A, Jaron KS, Kolarik M et al. (2016) White-nose syndrome without borders: *Pseudogymnoascus destructans* infection tolerated in Europe and Palearctic Asia but not in North America. Scientific Reports 6: 19829.

